# A sleep disturbance method by novel objects in the home cage to minimize stress

**DOI:** 10.1101/2024.08.26.609657

**Authors:** Christine Egebjerg, Mie Gunni Kolmos, Klas Abelson, Birgitte Rahbek Kornum

## Abstract

**Background:** The increasing prevalence of low sleep quality is a significant issue, particularly among adolescents, necessitating a deeper understanding of its biological consequences. In sleep research, various protocols are used for sleep deprivation or disturbance, each presenting its own set of confounding factors crucial to consider.

**New Method:** We developed a standardized seven-day sleep disturbance (SD) protocol using daily four-hour exposures to novel objects to minimize rodent stress. Objects were selected and characterized for wake-promoting properties, and exposure timing was structured to reduce variability and enhance experimental reliability and reproducibility.

**Results:** During the four hours of SD, the mice were efficiently sleep-deprived on the first and seventh day of SD. Thus, the selected objects efficiently sleep restricted the mice. On the first day of SD, the protocol induced sleep deprivation effect when measured over 24h, but by the seventh day, the mice recovered the sleep loss. Thus, this method is a sub-chronic sleep disturbance and not sleep deprivation. Fecal corticosterone concentrations remained unchanged during the seven days of SD.

**Comparison with existing methods:** This approach reduced the risk of stress through voluntary rather than forced wakefulness. Previously, novel objects have been exchanged randomly during mouse sleep initiation causing protocol variability and very frequent disturbances. Our protocol minimizes this by introducing the novel object in a structured manner.

**Conclusion:** We effectively disturbed the sleep of the mice during seven days without inflicting substantial stress. We further demonstrate the value of validating the efficiency of an SD protocol with 24h recordings.

**Highlights:**

- Standardized sleep disruption method by object exposure in mice
- Mice are sleep-restricted during the four-hour SD intervention all 7 days
- Mice habituate to a subchronic sleep deprivation setup by recovering sleep outside of the SD period
- Fecal corticosterone samples showed no difference before and after SD intervention

## 1 Introduction

Sleep research is a growing field, and sleep deprivation or disturbance is an often-used method to study the biological consequences of poor sleep. Rodent models are commonly used to investigate the molecular and cellular changes induced by lack of sleep (Bian et al., 2022; Colavito et al., 2013; Longordo et al., 2011; Murack et al., 2021; Tuan and Lee, 2019). However, no protocol has been established as the golden standard. It is difficult to keep a mouse awake without affecting other pathways than those of sleep, for instance, stress pathways (Nollet et al., 2020). This means that most protocols have confounding effects that must be considered when designing experiments.

The gentle handling protocol, where mice are kept awake by gently touching or stroking them or by shaking or tapping the cage, is commonly used (Colavito et al., 2013; Longordo et al., 2011; Murack et al., 2021). This method is easy to implement at a low cost, but it is highly dependent on the experimenter, increasing the risk of variability and decreasing reproducibility. Additionally, studies have reported elevated corticosterone concentrations levels in blood serum with this method (Longordo et al., 2011; Murack et al., 2021). One study investigated whether mice would habituate to gentle handling and whether corticosterone concentration levels in blood serum would increase after just 3 minutes of gentle handling in the light phase over 6 days. The results showed that the mice experienced a decrease in resting time after gentle handling on both day 1 and day 6. However, serum corticosterone concentration only increased at day 6, indicating sensitization of the stress axis. (Longordo et al., 2011).

A second method of sleep deprivation involves placing mice on small platforms over water. This method is primarily used for REM deprivation, as muscle atonia during REM sleep causes the mice to risk falling into the water so instead they wake up just as they enter REM sleep (Mendelson et al., 1974; Youngblood et al., 1997). Very small platforms have also been used for total sleep deprivation (Tuan and Lee, 2019). A confounding factor with this method is forced immobility. To control this, rodents can be placed on larger platforms where they can sleep but still experience forced immobility. Elevated serum corticosterone has been measured after the platforms over water method after a four-day setup (Suchecki et al., 1998; Youngblood et al., 1997) although conflicting results have been published after chronic REM deprivation (Arthaud et al., 2015; Khan et al., 2021; Suchecki et al., 1998; Youngblood et al., 1997). In addition, elevated corticosterone levels were measured in the control group placed on larger platforms over water in a four-day setup (Suchecki et al., 1998).

Another often-used approach is an automated setup (e.g. a moving beam or a jumping cage bottom) (Atrooz et al., 2022; Bian and De Lecea, 2023; Li et al., 2022). This approach standardizes the protocol but mechanically forces the animals to move and thereby keeps them awake. These systems can be set up in a closed loop with an EEG setup to track when sleep is initiated which will then elicit the mechanical wake stimuli (Bian and De Lecea, 2023; Hines et al., 2013). Elevated serum corticosterone levels have also been reported when using an automated setup (Huang et al., 2022; Yuan et al., 2021). One study shows that after 5 hours of sleep deprivation by a moving beam, young (10-14 week-old) and old mice (69-76 week-old) had increased plasma corticosterone (Yuan et al., 2021). It has been argued that this approach induces more movement than in the control mice, but that also depends on the specific setup (Bian and De Lecea, 2023). When in a closed-loop system, the mouse is only stimulated when falling asleep. This minimizes the induced excess movement. Systems that are not in a closed loop simply stimulate the mouse at random or set intervals and thereby risk inducing unwanted movement. In conclusion, all the mentioned methods have various issues, such as stress, forced immobility/mobility, or lack of standardization. An advanced setup using a closed-loop approach with EEG/EMG is necessary to circumvent some of these issues. However, the flexibility of a sleep deprivation experiment could be reduced if it is reliant on EEG/EMG equipment. Additionally, many research groups lack the capacity to implement such advanced setups, highlighting the need for a simple and easy sleep deprivation method. Furthermore, the diverse approaches to sleep deprivation make it challenging to compare results across experiments, emphasizing the need for standardized protocols.

When investigating sleep in adolescence, the possible activation of stress pathways becomes even more important to consider. Numerous studies have reported major consequences of stress exposure during adolescence, including long-term effects on emotional behaviors, social behaviors, and alterations in the hypothalamic-pituitary-adrenal (HPA) axis (Bekhbat et al., 2021; Cotella et al., 2019; McCormick et al., 2010; Rudolph, 2002). Cotella et al. demonstrated that chronic variable stress during adolescence has lasting effects in rats, increasing anxiety-like and despair-like behaviors, whereas similar stress exposure in adulthood did not result in long-term effects (Cotella et al., 2019). Consequently, when working with adolescent rodents, it is essential to choose a sleep deprivation method that ensures the measured effects are due to sleep deprivation rather than being confounded by stress.

Introducing novel objects into the home cage has been widely explored as an effective method for sleep disturbance (SD), as it promotes voluntary wakefulness and minimizes the stress associated with forced wakefulness. Although the introduction, timing and description of object is sparse calling for a protocol describing just that. Adolescent rodents, in particular, exhibit increased interest in novel objects and environments, making this method particularly useful for this age group (Adriani et al., 1998; Douglas et al., 2003). Many laboratories are already using novel objects to induce wakefulness, but the lack of standardization leads to poor reproducibility.

Here, we aim to create a standardized method using novel objects to induce SD. Our first objective was to identify objects that effectively promote wakefulness. Next, we developed and validated a standardized protocol for up to seven days of four-hour SD using novel objects.

## 2 Materials and methods

### 2.1 Animals

C57bl/6JTac female mice (purchased at Taconic) were used for these experiments with access to ad libitum food and water. The lighting conditions were 12:12 h light: dark cycle and temperature and humidity were kept constant. All mice were group housed (2-5 mice per cage) with enrichment such as nesting material, wooden blocks, a hemp robe hanging from the lid, a plastic tube, and a plastic hide when not video recorded, EEG/EMG recorded or habituated in EEG cages. In the recording setup, all the enrichment tools were the same except the plastic tube and hide, which were excluded due to complications with the cord from the head mount. The objects are visible to the experimenter hence blinding is not possible for this setup. The protocol was approved by the Danish Animal Experimental Inspectorate (license number: 2022-15-0201- 01194). Per the EU Directive 2010/63/EU on the protection of animals used for scientific purposes, mice were monitored for signs of compromised health or the loss of more than 20% of their body weight.

### 2.2 Selection of Objects

To select objects and determine the timing and setup of object exposure for SD, eight female C57BL/6Rj mice were single-housed in EEG cages (9000-K20, Pinnacle Technology Inc.) equipped with infra-red video cameras (9000-K10, Pinnacle Technology Inc.) and habituated for three days before testing.

21 objects were tested by exposing mice to one single object for 30 minutes at a time and manually scoring for how long each mouse interacted with the object (Figure 1A). The objects were selected to be easily accessible by being available in a pet shop or from regular lab equipment. Objects with a total interaction time of less than five minutes were discarded to select the most effective wake-promoting objects. Of the remaining objects, only objects with at least one interaction bout longer than 30 seconds were selected for the final study (Figure 1B). Sleep/wake behavior was based on observation during the object exposure, here sleep was defined as the mouse being immobile and resting in the nest.

**Figure 1:**
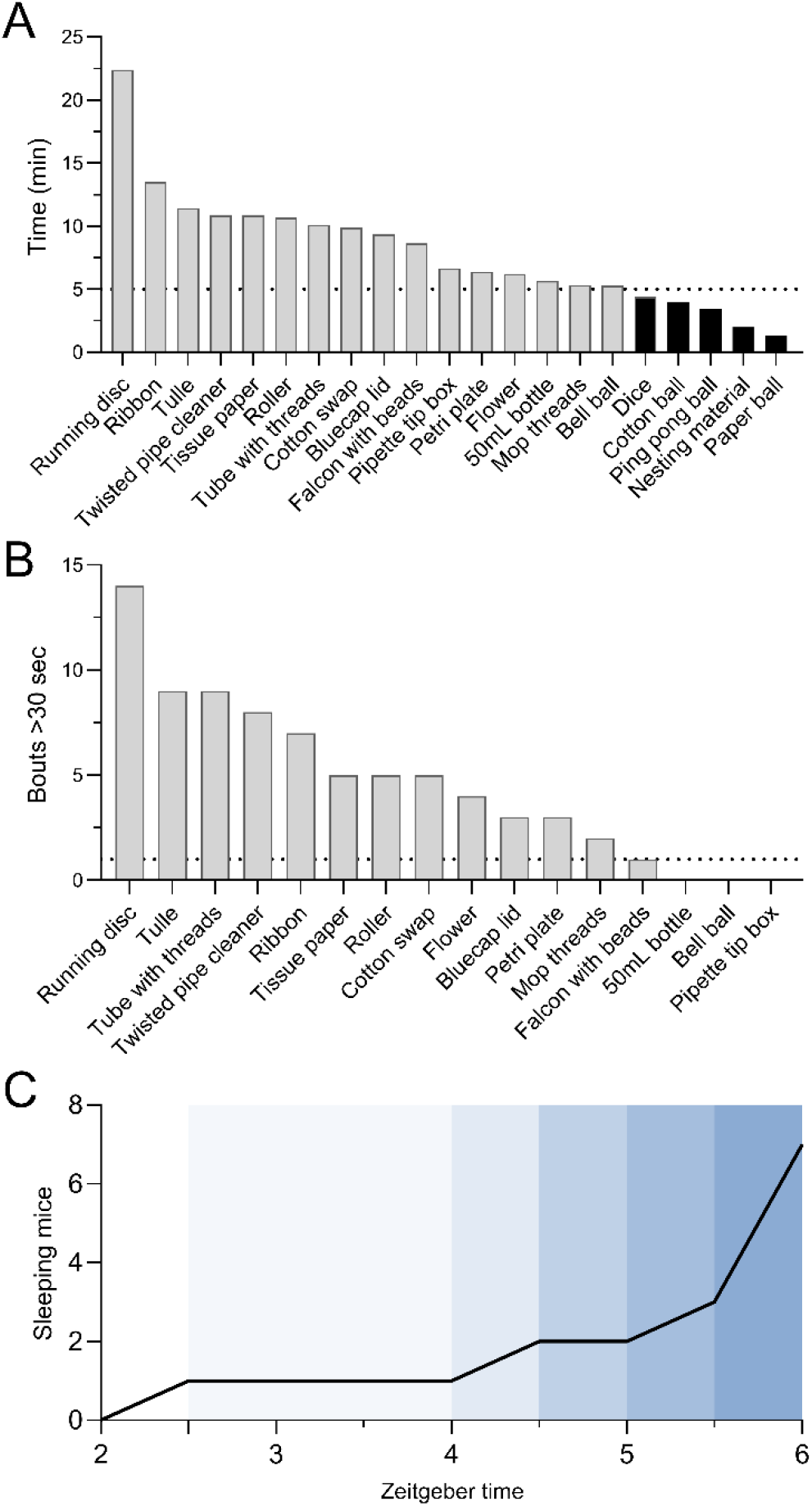
Testing capability of objects to keep mice engaged for sleep deprivation. A) Total interaction time in minutes with objects during a test time of 30 minutes for each object. The horizontal dotted line indicates the cut-off value of 5 minutes. Objects with a total interaction time below 5 minutes were excluded (indicated by black bars). B) Number of interaction bouts that were above 30 seconds during the 30-minute test time. The horizontal dotted line indicates the cut-off value of one bout. Objects that did not sustain interaction for more than 30 seconds at least once during the test were excluded. n=1, each object was tested on a single mouse, eight mice were used in total to obtain simultaneous testing. C) The remaining 13 objects were tested again in a 4-hour sleep deprivation, rotating the objects between eight single-housed mice every 30 minutes. The graph indicates the number of mice sleeping (by visual assessment) at any time during the preceding 30 minutes. n=1 was tested once on eight mice. Colored shading indicates the number of mice sleeping.

### 2.3. Protocol design

For the final protocol of SD by novel object exposure, the 13 objects (Fig. 1A and 1B, Suppl. Table 1) that passed the aforementioned interaction time and bout number criteria were systematically alternated between cages. To maintain novelty, no object (except for the running disc) was introduced two days in a row during SD. The investigated objects were categorized into high (running disc), medium (e.g., tulle), and low (e.g., flower) effects, and an exposure scheme was made to ensure that two low-effect objects did not occur subsequently. Running discs were selected instead of running wheels to enable simultaneous EEG/EMG recording. The administration of the objects was done with minimum disruption of the cage. They were placed in the cage without touching the mouse, and if the object became integrated into the nest, it was gently removed without ruining the nest. Most objects were not cleaned between introductions to different cages during the protocol for practical reasons although the paper towels were discarded after use due to deterioration. After the four-hour use, the objects of plastic were cleaned with ethanol, and textured objects were cleaned by picking out the wood chips and fecal boli. If the objects were too damaged, they were discarded. To standardize the odor of the objects, new objects were therefore put in a non-experimental cage with mice of the same sex prior to exposure to ensure murine odor on all objects.

An example of a day of SD is attached in the supplements along with a list of objects (Suppl. Table 1 and 2). Since the running disc had the longest interaction time (Figure 1A) it was introduced in the SD protocol twice every day.

### 2.4 Validation of SD by EEG/EMG/video recordings

To validate the wake promotion of the objects, 8 female adolescent mice (SD start at p36-p42) were used for EEG/EMG recording and sampling of fecal boils for corticosterone measurements (Fig 2A). EEG/EMG surgery was performed 8 days prior to the SD start at p28.

**Figure 2:**
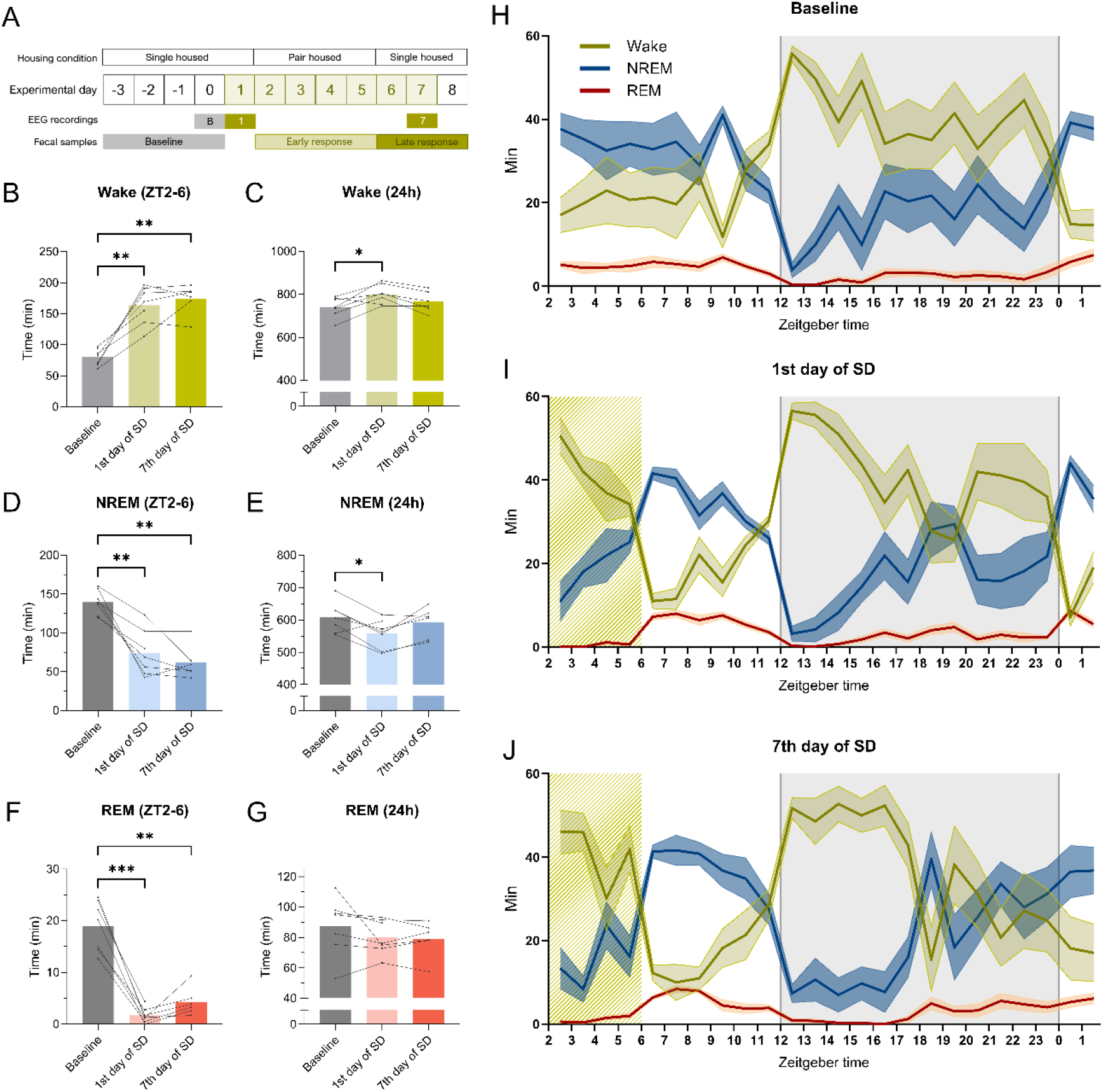
SD by novel object prolongs wakefulness and decreases the duration of NREM and REM but the sleep is recovered over the 24 hours on the seventh day of SD. Mice (n=8) had EEG/EMG/video recording on the baseline, first day of SD, and seventh day of SD to validate the SD by novel objects. A) Experimental overview of the EEG experiment and fecal sampling B) Duration of wakefulness at ZT2-6 on all days. C) Total duration of wakefulness on all days. D) Duration of NREM at ZT2-6 on all days. E) Total duration of NREM on all days. F) Duration of REM at ZT2-6 on all days. G) Total duration of REM on all days. H) Duration of wake, NREM, and REM at baseline plotted per hour. The shaded gray indicates the dark phase (ZT12-0). The colored shading around the data indicates SEM. I) Duration of wake, NREM, and REM on the first day of SD plotted per hour. The yellow-striped area indicates exposure to novel objects. J) Duration of wake, NREM, and REM on the seventh day of SD plotted per hour. Data from the same mouse from timepoint to timepoint is connected by a line on bar graphs. Mixed-effect analysis, with the Geisser-Greenhouse correction and Tukey’s multiple comparisons tests, was used to analyze the data illustrated in the bar plots, *p < 0.05, **p<0.01, *** p<0.001.

#### 2.4.1 EEG/EMG surgery

All mice were anesthetized with isoflurane (induction 4%; 4L/min O_2_, maintenance 1.5-2.5; 2L/min O2) to start the surgery. After induction, the mice received analgesia (buprenorphine, 0.05 mg/kg, Bupaq Multidose, Nomeco, Denmark) and antibiotic drugs (Baytril, 5 mg/kg, Nomeco, Denmark), subcutaneously. The core temperature of the mice was monitored by a rectal probe and stabilized by a heating pad. Before attaching the mouse head to the stereotactic frame, the head and neck were shaved. When attached to the frame ophthalmic ointment was applied and the head and neck were cleaned with water and povidone-iodine. Local analgesia (lidocaine, Lidor Vet; Nomeco, Denmark) was injected subcutaneously at the site of the incision and subsequently, the incision (∼1cm) was made from the crown of the head to the neck. The skull was cleaned and dried with ethanol to prepare the attachment of the electrode. The frontoparietal bipolar 3-channel EEG/EMG head mount (#8201; Pinnacle Tech. Inc., KS, USA) was placed into a clamp on a standard probe holder to the stereotaxic frame to place and glued 3 mm anterior of bregma. Two craniotomies were drilled above the frontal lobe and two above the parietal lobe. Subsequently, two 0.10 mm screws were secured in the frontal holes and two 0.12 were secured in the parietal holes fitting in the prefabricated head mount. Screws were secured without penetrating the dura. Epoxy was carefully placed around the screws (#8226, Pinnacle Tech. Inc. KS, USA). EMG wires were placed intramuscularly bilaterally in the nuchal muscles. Dental cement was applied on the head mount to secure it, and sutures were applied in the neck if there was excess wound. To finish the surgery, a long-lasting analgesic (Mexilocam, 5mg/kg, Metacam, Nomeco Denmark) was injected subcutaneously and the mouse was placed in a cage with a heating lamp for recovery. Post-surgical care was performed three days after the surgery, which included weighing, administration of long-lasting analgesics, and a behavioral assessment.

#### 2.4.2 EEG/EMG recording and scoring

Habituation started three days before the baseline recording by housing the mice individually in a 10” circular transparent recording cage (#9000-K20; Pinnacle Tech. Inc., KS, USA). The mice were exposed to novel cages for the first 24 hours of habitation and subsequently attached to the cable connecting the EEG head mount to the EEG equipment. Three 24-hour recordings were performed - baseline, first day of SD, and seventh day of SD. Mice returned to home cages in between recordings on the first and seventh day of SD and re-habituated to EEG cages and cables one day prior to the seventh day of SD (Figure 2A). One mouse where euthanized between the first day and the seventh of SD due to a loose head mount reading to an n of 7 for the last recording.

To acquire the EEG/EMG and video recording, the Sirenia Acquisition software 2.2.12 (Pinnacle Tech. Inc. KS, USA) was used. The data was digitized at 400 Hz and processed through a preamplifier to filter out noise (×100 gain, high pass filtered at 0.5 Hz EEG and 10 Hz EMG). When scoring the data, it was further amplified (x50) and filtered (0.5 Hz EEG, 10 Hz EMG). The EEG/EMG signals were divided into four-sec epochs and wake, NREM, and REM were determined manually using standard criteria (Mang and Franken, 2012).

### 2.5 Fecal corticosterone measurements

During the experimental paradigm, fecal boli were sampled from EEG or home cages bedding material at three-time points – before SD, early SD, and late SD – to measure corticosterone (as immunoreactive fecal corticosterone metabolites [FCM]) in a non-invasive manner (Fig. 2A). Before SD, fecal boli were sampled from EEG cages just before novel object exposure on the first day. Early SD fecal boli were sampled from home cages when returning mice into EEG cages for the last recording. Late SD fecal boli were sampled from EEG cages after the last recording. The early SD timepoint only has only four samples due to double housing. All samples were collected at ZT2.

FCM was extracted from the fecal boli and quantified by ELISA, as previously described (Abelson et al., 2016; Falkenberg et al., 2019; Kalliokoski et al., 2015; Nørgaard et al., 2018). In brief, each fecal sample was weighed and submerged in 96% ethanol (3 ml/g feces), followed by incubation on a shaking table overnight for 12 h. The homogenate was centrifuged for 20 min at 3000 × g (Scanspeed 1236R, Labogen, Lynge, Denmark), whereafter the supernatant was decanted, and the pellet discarded. One ml of the supernatant was centrifuged for 15 min at 10,000 × g in a tabletop centrifuge (model 5415D, Eppendorf, Hamburg, Germany). A 300-μL aliquot of the supernatant was recovered by using a pipette and evaporated to dryness using an evaporator (model EZ2, Genevac, Ipswich, United Kingdom). The evaporate was then resuspended in 300 μL assay buffer and analyzed by using a competitive corticosterone ELISA (EIA-4164, DRG Diagnostics, Marburg, Germany) according to the manufacturer’s instructions. Cross-reactivities for the assay were reported as follows: Progesterone, 7.4%; deoxycorticosterone, 3.4%; 11- dehydrocorticosterone, 1.6%; cortisol, 0.3%; pregnenolone, 0.3%; and other steroids, <0.1%.

### 2.6 Statistics

To perform statistical analysis and plotting data, GraphPad Prism v10.2.3 (San Diego, CA, USA) was used. Mixed-effect analysis, with the Geisser-Greenhouse correction and Tukey’s multiple comparisons, was used for the paired data. An ordinary two-way ANOVA with main effects only and a Tukey’s multiple comparison test, with a single pooled variance was used to analyze the distribution count of the bout lengths (timepoint x count of bout length). For analyzing corticosterone concentration an ordinary one-way ANOVA test and a Holm-Šídák multiple comparisons test were used. Significant ANOVAs are reported with p-values and p-values < 0.05 were considered statistically significant and noted as *p<0.05, **p < 0.01, ***p < 0.001 on the figures. P-values between 0.1-0.05 are noted with the specific p-value in the figures.

## 3 Results

### 3.1 Selection of objects

Testing novel objects for wake-promoting effects resulted in 13 useful objects. The running disc was the most effective (Fig. 1A and 1B). To estimate the efficacy of the object exposure, it was manually noted how many mice were sleeping in their nests at any time during 30 minutes with an object during SD testing for four hours from ZT2-6 (Figure 1C). This showed that the sleep pressure increased already 2 hours into SD, and that novel objects were no longer able to keep any of the 8 mice awake after four hours of exposure (Fig. 1C). These results show that not all objects are useful for SD, especially as sleep pressure increases during the procedure. It is therefore important to always validate objects before using them for SD.

### 3.2 Validation of SD by EEG/EMG and examination of changes in the sleep architecture

Next, we investigated how the SD by novel objects affected sleep using EEG/EMG. Wake, NREM, and REM duration per hour at baseline, the first day of SD, and the seventh day of SD showed a different pattern during the 24-hour recording (Fig. 2H-J). Between ZT2-6, when the mice received the novel objects, the wake duration was increased on the first and seventh day of SD compared to baseline (Fig. 2B, p <0.01). Duration of NREM and REM sleep decreased during SD compared to baseline (Fig. 2D and 2F, p<0.01, p<0.001). For the total 24 hours, the wake duration increased on the first day of SD compared to baseline but not for the seventh day of SD (Fig. 2C, p<0.05). NREM duration decreased on the first day of SD but showed no difference between the baseline and the seventh day (Fig. 2E, p<0.05). There were no changes in the duration of REM sleep between the time points for the 24-hour recording. These results show that the mice are sleep deprived during the four-hour SD both on the first day of SD and after a week of daily SD. Further, when accounting for the total 24-hour recording, the mice were only deprived of NREM sleep on the first day but not on the seventh day of SD where time in all vigilance states were similar to baseline.

To understand the fragmentation of sleep, bout counts and durations of the vigilance states were analyzed. During the SD (ZT2-6), the mean wake bout count decreased on the seventh day of SD (Fig. 3A, p < 0.05). There was no significant change on the first day of SD, but a tendency for reduction of mean wake bouts (Fig. 3A, p = 0.0579). The mean duration of wake bouts increased on both experimental days compared to baseline (Fig. 3B, p < 0.01). NREM and REM bouts decreased in number on both experimental days (NREM: Fig. 3C, p < 0.05, p < 0.01 and REM: 3E, p < 0.01, p < 0.001) but had the same bout duration as at baseline (Fig. 3D and 3F). These results indicate that the four-hour SD increased wake bout duration and decreased the number of NREM and REM bouts yet did not fragment sleep.

**Figure 3:**
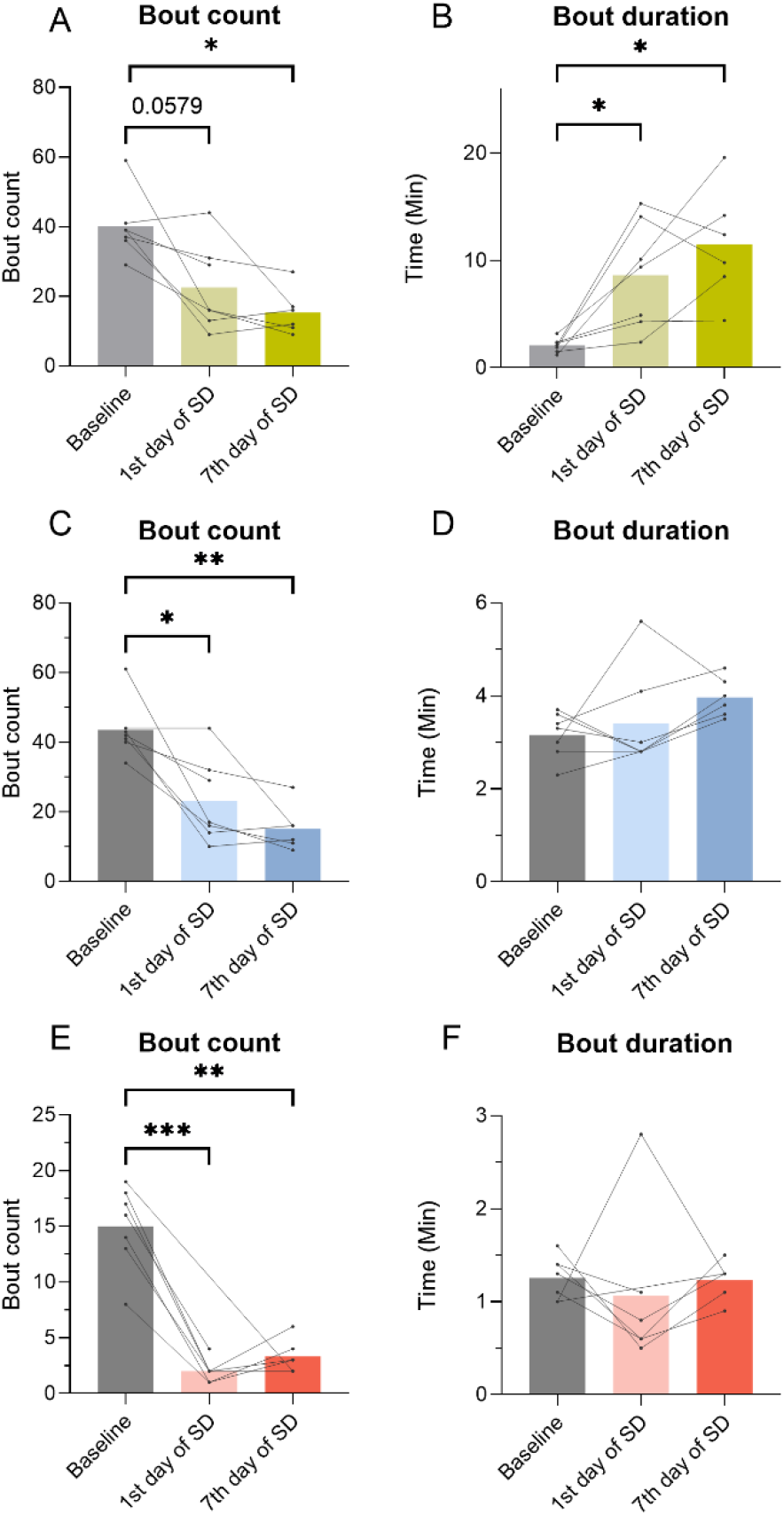
The number of all vigilance state bouts is reduced whereas wake bout duration increases during SD (ZT2-6). Bout count and duration of wake, NREM, and REM were calculated from the mice (n=8) during SD (ZT2-6) at all time points. A) Wake bout count. B) Wake bout duration. C) NREM bout count. D) NREM bout duration. E) REM bout count. F) REM bout duration. Data from the same mouse from timepoint to timepoint is connected by a line. Mixed-effect analysis, with the Geisser-Greenhouse correction and Tukey’s multiple comparisons tests, was used to analyze the data illustrated in the bar plots, *p < 0.05, **p<0.01, *** p<0.001.

During the 24-hour recording, bout counts for the vigilance states were unchanged between all time points (Fig. 4A, 4D, and 4G). The bout duration of wake increased on the first day of SD but not on the seventh day of SD (Fig. 4B, p< 0.05). There was no significant change in bout duration for NREM sleep (Fig. 4E). The duration of REM bouts increased on the seventh day of SD when compared to baseline and the first day of SD (Fig. 4H, p <0.05). The distribution of bout counts was analyzed by using the percentage distribution of each bout count to accommodate for changes in group size. There were no changes observed in the distribution of the bout for any of the vigilance states (Fig. 4C, 4F, and 4I). These results indicate that the mean NREM and REM sleep bouts became longer on the seventh day of SD.

**Figure 4:**
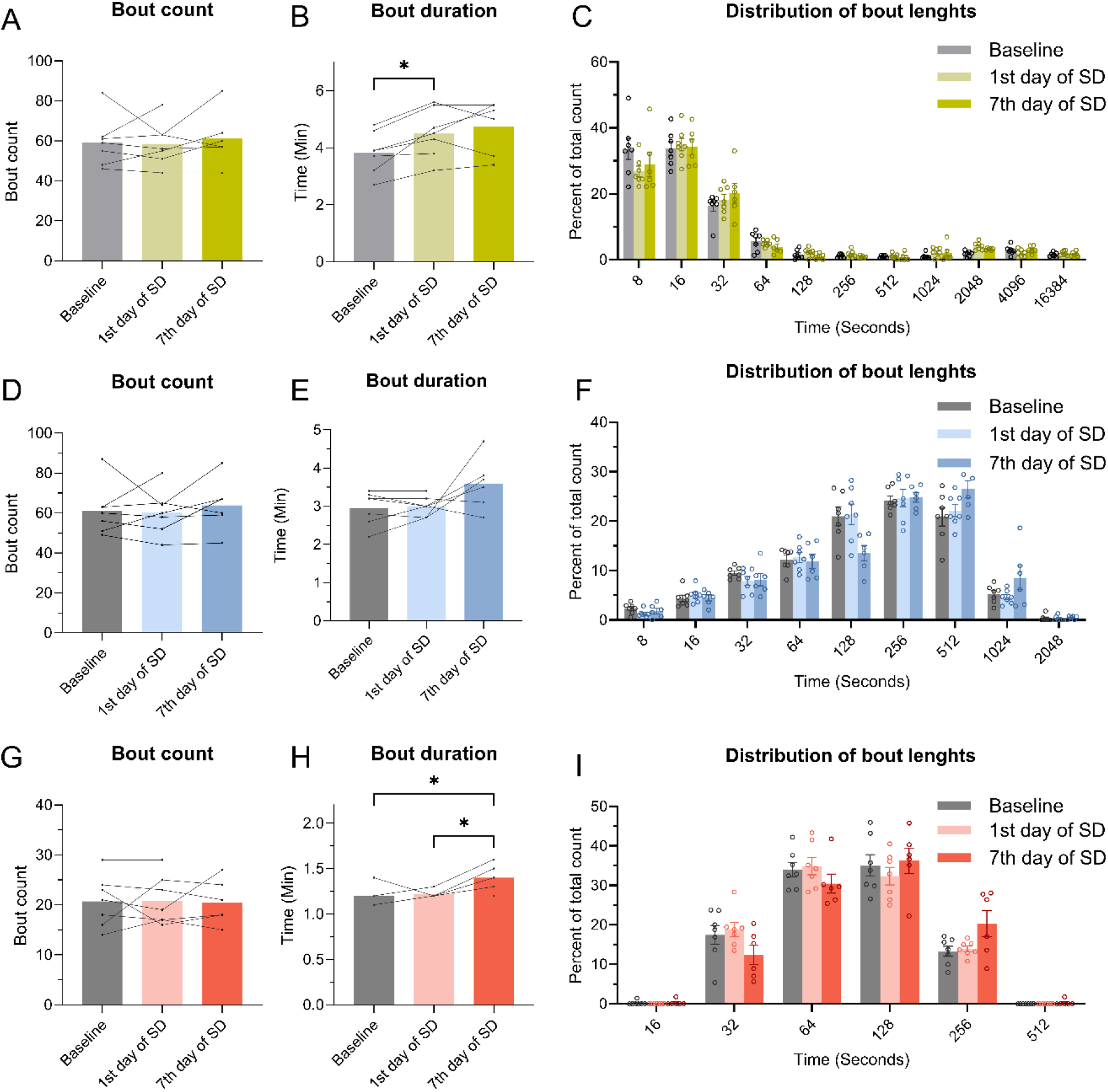
Over the total 24 hours, the bout count is unchanged, but all vigilance states show indications of longer bout duration. The bout counts and duration of wake, NREM, and REM were calculated from the EEG data (n=8) during 24 hours on all days. A) Wake bout count. B) Wake bout duration. C) Distribution of bout lengths of the wake bouts in the percentage of the total count. D) NREM bout count. E) NREM bout duration. F) Distribution bout lengths of the NREM bouts in the percentage of the total count. G) REM bout count. H) REM bout duration. I) Distribution of bout lengths of the REM bouts in the percentage of the total count. For A, B, D, E, G, and H mixed-effect analysis, the Geisser-Greenhouse correction and Tukey’s multiple comparisons test were used to analyze the data, *p < 0.05. For C, F, and I an ordinary two-way ANOVA with main effects only and a Tukey’s multiple comparison test, with a single pooled variance was used to analyze the data, *p< 0.05, *p<0.01.

### 3.3 Validation of stressless protocol

To test whether the mice were stressed during the SD, fecal samples were collected, and the concentration of corticosterone (as immunoreactive fecal corticosterone metabolites) was determined. There was no significant change in the concentration of corticosterone between baseline, early SD, and late SD (Fig. 5). This indicates that the mice were not substantially stressed by the daily exposure to novel objects.

**Figure 5:**
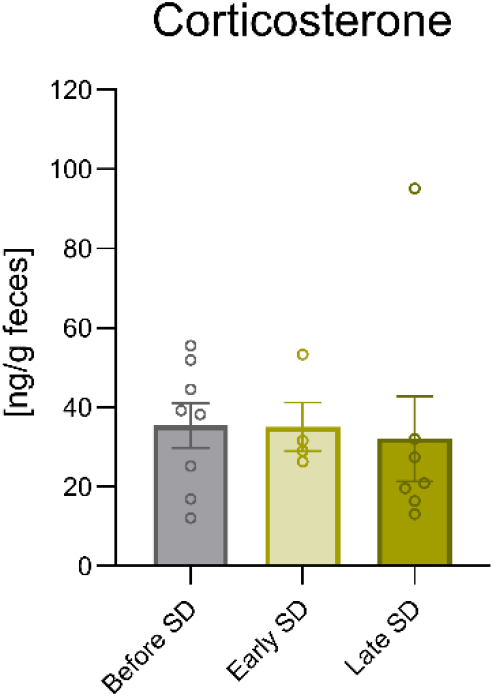
There was no change in the concentration of corticosterone between the baseline and the SD days. Fecal boli from the mice were sampled over multiple days. Before and the late SD had a sample size of 8 as the mice were single house. At early SD they were double housed thus having a sample size of 4. An ordinary ANOVA test and a Holm-Šídák multiple comparisons test were used. The bar plot is illustrated with mean ± standard error of the mean (SEM).

## 4 Discussion

The goal of this study was to develop a less stressful SD model using the intrinsic exploratory behavior of the mice, hence inducing voluntary wakefulness. First, objects were selected based on their wake-promoting abilities, and a sequence of introduction of objects was developed. Next, the protocol was validated by sleep quantification using EEG/EMG. Our SD protocol resulted in an increased duration of wakefulness during the introduction of objects (ZT2-6), showing an effective SD method both acutely on the first day and after seven days of SD. Over the 24 hours, the mice were also sleep deprived on the first day of SD, but on the seventh day, the mice recovered their lost sleep at other time points, suggesting habituation to the setup. We, therefore, demonstrate that novel objects can be used to sleep-deprive mice during ZT2-6 for seven days in a row, but this is not a method for total sleep deprivation beyond day 1.

Due to the voluntary nature of our protocol, mice did sometimes sleep during object exposure. In these instances, the NREM and REM bouts had the same duration as during baseline, but the mice experienced fewer bouts during SD. Outside SD the mean REM bout duration increased, which could be observed in the total 24-hour recording.

For EEG/EMG recordings mice undergo surgery to implant the EEG/EMG electrode. The mice carry this implant together with a preamplifier attached to the EEG/EMG recording system during the recordings. The weight on their heads might reduce the mice’s interest in interacting with objects. Hence the novel object might work more efficiently on a non-EEG/EMG mouse although this would have to be validated in a future setup during the seven days of SD.

Since this method is only sleep-depriving mice on the first day when considering all 24 hours, we define our protocol as a paradigm of sleep disturbance (SD) rather than sleep deprivation. It is important to consider all 24 hours when performing a sub-chronic SD study as sleep recovery may occur. Most studies only validate the SD paradigm during the SD period, the first 24 hours of SD, or not at all, but do not take into consideration that the mice will habituate and catch up on sleep outside the SD time window (Atrooz et al., 2022; Bian et al., 2022; Murack et al., 2021; Tuan and Lee, 2019). If the goal is to study the consequences of sleep loss and not disturbance, certain protocols will not be the right ones to choose. The phenomena of sleep rebound outside observation times is also important to consider when interpreting findings from SD protocols in the literature.

The protocol we describe here does not induce elevated corticosterone levels in fecal samples. We chose to perform fecal sampling to not induce injection stress to the adolescent mice thus exact measurement after administration of objects are difficult to assess. This suggests that the mice are not stressed by our SD protocol, in contrast to what has been seen with a gentle handling protocol (Longordo et al., 2011; Murack et al., 2021), platforms over water (Mendelson et al., 1974; Youngblood et al., 1997), and automated setups (Huang et al., 2022; Yuan et al., 2021). One study also showed increase serum corticosterone immediately after the gentle handling protocol, but the mice showed decreased corticosterone on the sixth’s day. This indicates mice could potentially habituate to a gentle handling protocol although forceful matter of the protocol still persists.

An increased concentration of corticosterone in blood plasma, urine, or fecal samples is often used as a measure of stress in rodents (Abelson et al., 2016; Kamakura et al., 2016; Kim et al., 2013). Cortisol in humans, the equivalent to corticosterone in rodents, is also associated with increased stress and anxiety (Elnazer and Baldwin, 2014; Hellhammer et al., 2009). However, cortisol has a complex effect in relation to mood (Shirtcliff et al., 2014). Studies have shown that an increase in cortisol is not only associated with negative emotions related to stress but also alertness, activeness, and excitement (Hoyt et al., 2016; Mazur et al., 1997). Similarly, mice show increased corticosterone when exposed to an enriched environment (Moncek et al., 2004). Thus, corticosterone measurements are difficult to interpret as just a measure of negative stress.

It can be argued that our protocol uses short enrichment to generate wakefulness. Additionally, the running disc could also increase the locomotion activity which may influence the outcome of an SD experiment. Environmental enrichment has been shown to decrease the effect of the stress paradigms and thus enrichment with novel objects could potentially interfere with the effect of SD (Benaroya-Milshtein et al., 2004). Since both SD and enrichment previously have been shown to increase corticosterone, and we observe no changes, we can argue that the mice are neither affected by the stress of SD nor enrichment from the novel object. In general, separating sleep deprivation from stress effects is a challenge in rodent models (Nollet et al., 2020), and important to consider when interpreting findings from different SD protocols. Potentially more important than elevated corticosterone levels, is the consideration of the translational value of the selected protocol. When considering the brain pathways involved in SD a common feature would probably be the activation of HPA, but the upstream brain activation could vary and potentially cause different consequences of SD. Not only the stress pathway can lead to extended wakefulness, but also factors such as an uncomfortable environment (Mori et al., 2021), fear (Pawlyk et al., 2005; Werner et al., 2020), and arousal (Dresp-Langley and Hutt, 2022). The protocol presented here aims to model human activities where the brain is positively aroused without a stressful experience.

The protocol we describe here is particularly useful for studying the consequences of disrupted sleep during adolescence, due to the naturally high level of exploratory behavior in this age group. It is also a simple and low-cost method that is easy to implement. For implementation, we recommend using the object suggested here or validating new wake-promoting objects by characterizing interaction time and interaction bout lengths. We do not recommend using this for more than 4 hours SD as the sleep pressure will then exceed the urge to explore novelty.

## 5. Conclusion

Here we developed a standardized, easy, low-cost SD protocol without a detectable stress response in the mice. First, wake-promoting objects were characterized, and a protocol was developed. Next, the protocol was validated by EEG/EMG. The mice were sleep-deprived during the introduction of objects (ZT2-6) and on the first day of SD. On the seventh day of SD, the sleep deprivation effect was lost. We also show data indicating that sleep is less fragmented after SD. During the setup, fecal samples were collected for corticosterone measurements and there were no changes between timepoints. For a sub-chronic sleep deprivation protocol, it is crucial to validate sleep deprivation during the setup and not only on the first day. The mice might show compensatory rebound sleep and therefore not be fully sleep deprived. The protocol we present here is a sleep disturbance protocol especially suited for examining the effects of disturbed sleep during adolescence.

## Declaration of interest: None

Declaration of generative AI and AI-assisted technologies in the writing process During the preparation of this work the author(s) used ChatGPT in order to improve the readability of the text, and correct typos and grammar. After using this tool/service, the author(s) reviewed and edited the content as needed and take(s) full responsibility for the content of the publication.

## Abbreviations

EEG –: electroencephalography
EMG –: electromyography
FCM –: fecal corticosterone metabolites
NREM –: non-rapid eye movement
REM –: rapid eye movement
SD –: sleep disturbance
ZT –: zeitgeber time

## Acknowledgments

Lundbeck Foundation for funding the project,

## Supplementary

**Supplementary table 1:**
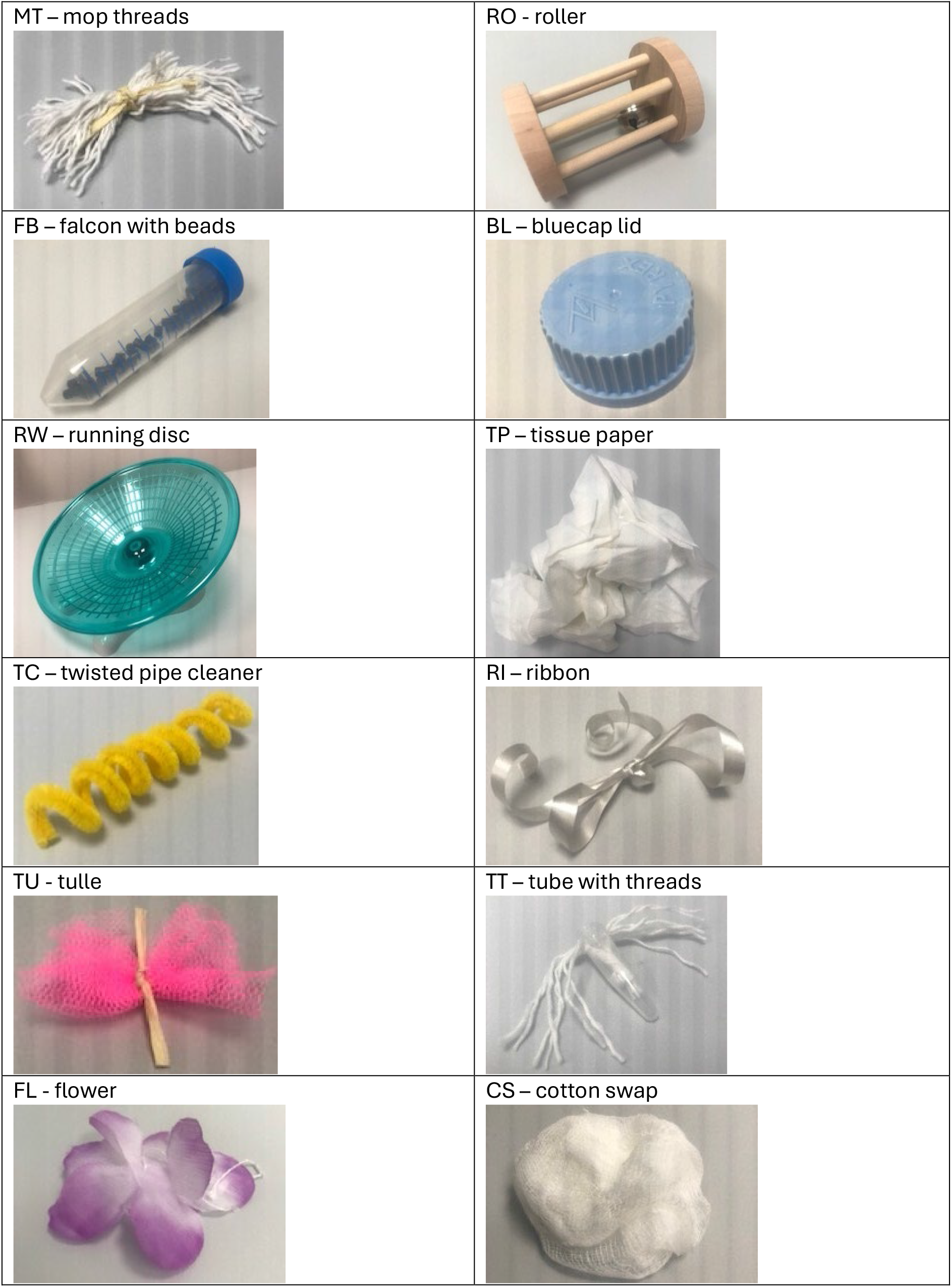
Overview of object selected for wake promotion.

**Supplementary table 2:**
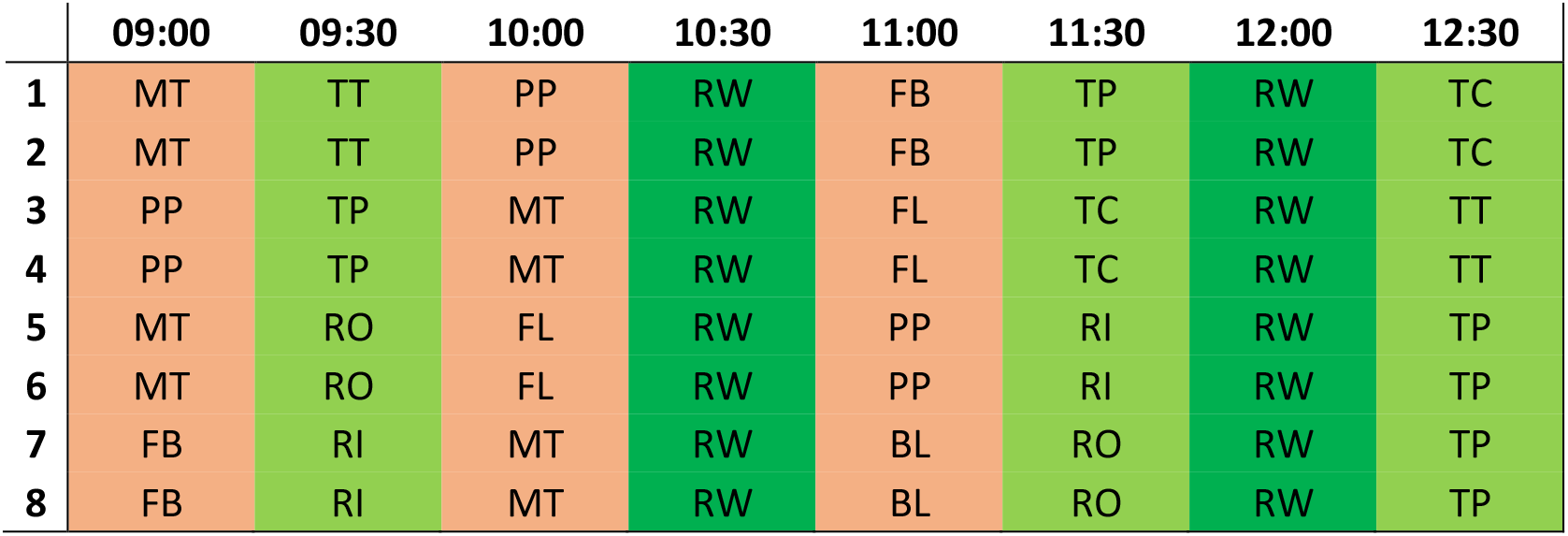
Example of an SD day. Red coloring indicates low interaction, light green indicates medium interaction and dark green indicates high interaction.

