## Supplemental figures for "A sleep disturbance method by novel objects in the home cage to minimize stress"

**Supplementary**

| | |
|---|---|
| MT – mop threads 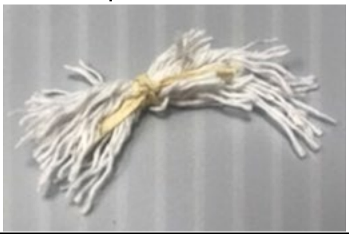 | RO - roller 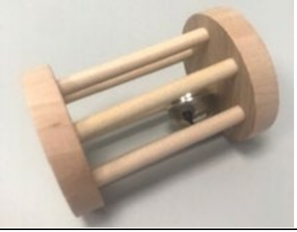 |
| FB – falcon with beads 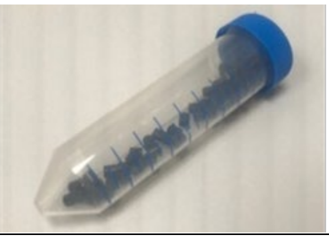 | BL – bluecap lid 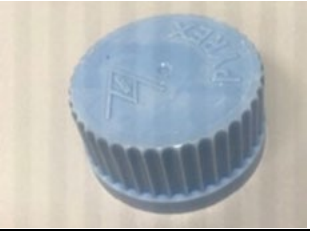 |
| RW – running disc 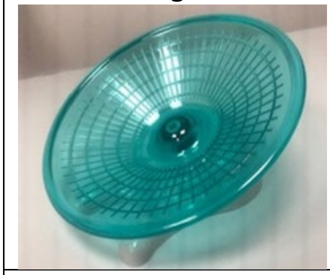 | TP – tissue paper 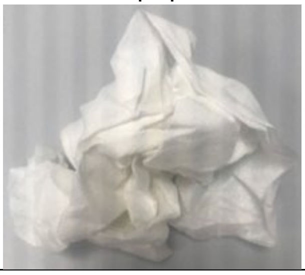 |
| TC – twisted pipe cleaner 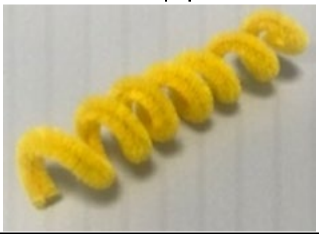 | RI – ribbon 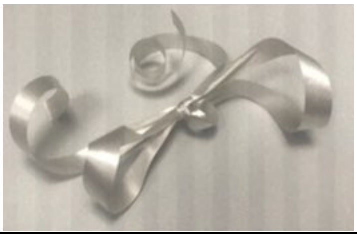 |
| TU - tulle 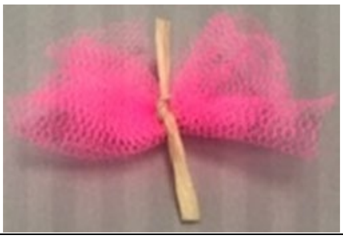 | TT – tube with threads 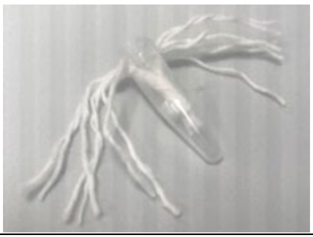 |
| FL - flower 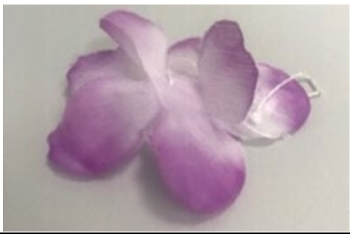 | CS – cotton swap 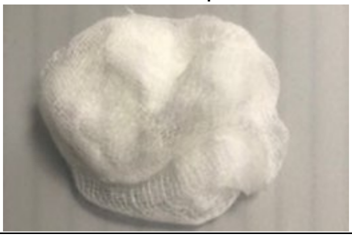 |

**Supplementary table 1: Overview of object selected for wake promotion.**

| | 09:00 | 09:30 | 10:00 | 10:30 | 11:00 | 11:30 | 12:00 | 12:30 |
|---|---|---|---|---|---|---|---|---|
| **1** | MT | TT | PP | RW | FB | TP | RW | TC |
| **2** | MT | TT | PP | RW | FB | TP | RW | TC |
| **3** | PP | TP | MT | RW | FL | TC | RW | TT |
| **4** | PP | TP | MT | RW | FL | TC | RW | TT |
| **5** | MT | RO | FL | RW | PP | RI | RW | TP |
| **6** | MT | RO | FL | RW | PP | RI | RW | TP |
| **7** | FB | RI | MT | RW | BL | RO | RW | TP |
| **8** | FB | RI | MT | RW | BL | RO | RW | TP |

**Supplementary table 2: Example of an SD day.** Red coloring indicates low interaction, light green indicates medium interaction and dark green indicates high interaction.
